# The presence of BBB hastens neuronal differentiation of cerebral organoids – the potential role of endothelial derived BDNF

**DOI:** 10.1101/2022.07.22.501119

**Authors:** Giorgia Fedele, Alessandra Cazzaniga, Sara Castiglioni, Laura Locatelli, Antonella Tosoni, Manuela Nebuloni, Jeanette A. M. Maier

## Abstract

Despite remaining the best in vitro model to resemble the human brain, a weakness of human cerebral organoids is the lack of the endothelial component that in vivo organizes in the blood brain barrier (BBB). Since the BBB is crucial to control the microenvironment of the nervous system, this study proposes a co-culture BBB and cerebral organoids. We utilized a BBB model consisting of primary brain microvascular endothelial cells and astrocytes in a transwell system. Starting from induced Pluripotent Stem Cells (iPSCs) we generated human cerebral organoids which were then cultured in the absence or presence of an in vitro model of BBB. We evaluated if the presence of the BBB influences the maturation of cerebral organoids. By morphological analysis, it emerges that in the presence of the BBB the cerebral organoids are better organized than controls in the absence of the BBB. This effect seems to be driven by Brain Derived Neurotrophic Factor (BDNF), a neurotrophic factor released by the endothelial component of the BBB, which is involved in neurodevelopment, neuroplasticity and neurosurvival.

**Graphical Abstract:** 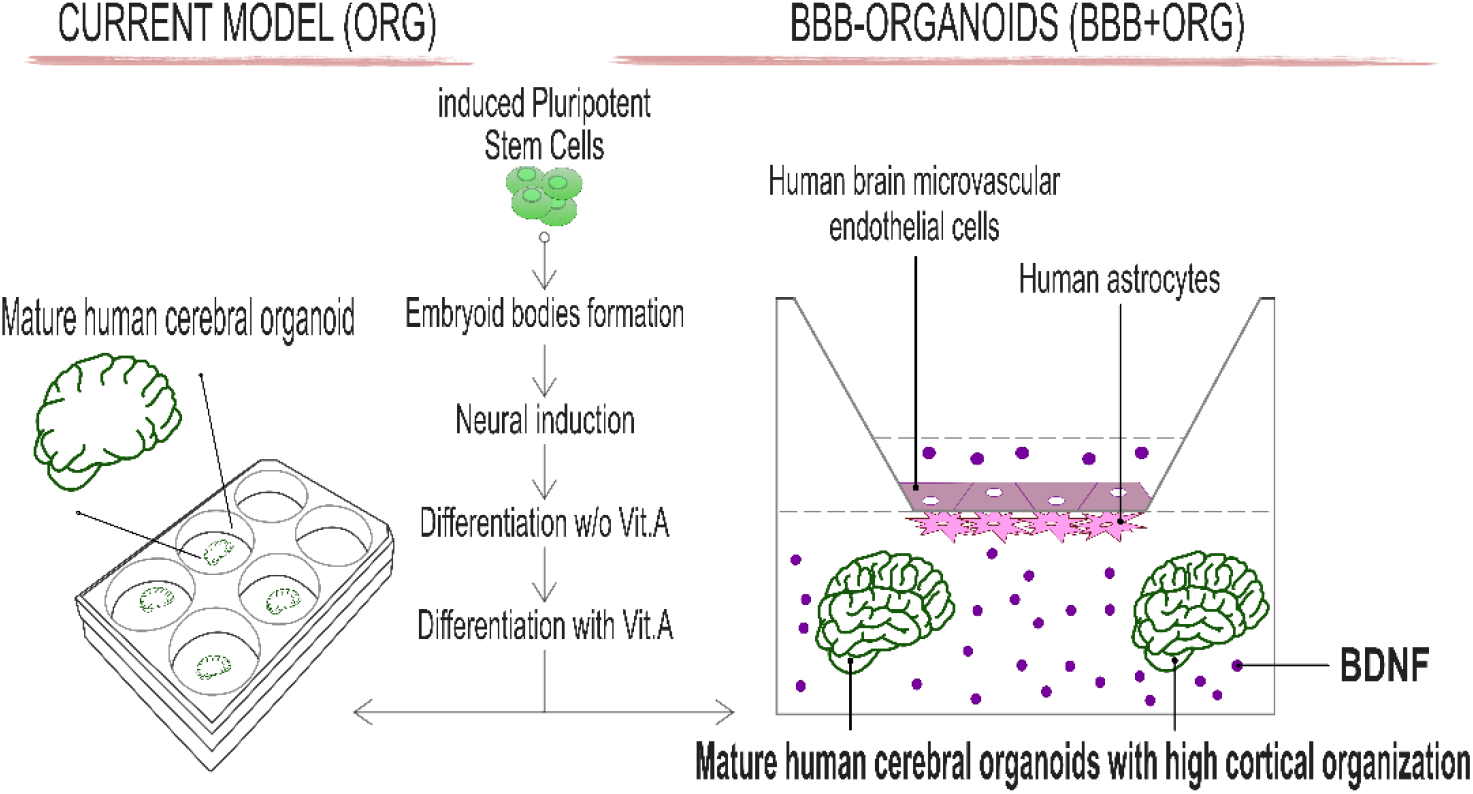

The current culture model of human cerebral organoids does not require the presence of a BBB (left side). However, the BBB is an important source of BDNF, which is crucial for neurodevelopment and brain health. The cerebral organoids co-cultured for 4 days in the presence of the BBB show a higher cortical organization than the organoids cultured in the absence of the BBB, as illustrated on the right.

## 1. Introduction

Human brain is an extraordinarily unique and complex structure that controls actions and behaviour, interprets senses, handles complicated cognitive tasks, stores memories, elaborates feelings and governs all the processes that regulate the body. These highly integrated functions rely on a significant cellular and architectural diversity. Although far from being complete, our present knowledge on how the brain functions derives from experimental models that can only mimic some aspects of brain complexity. The limits of classic 2D cultures of nervous cells are primarily due to the homogeneity of the neural cultures. Experiments performed on animals, mainly rodents, while contributing to the understanding of the role of some genetic mutations in neurogenesis [1], were seldom translated to clinical setting. On the other hand, it is well known that rodents and humans are evolutionarily very distant and human brain is much more complex than rodents’, in particular when focusing on cortical folding patterns [2]. In mice 43 different areas per each hemisphere were identified in face of the 180 human cortical areas, and this finding is in agreement with the increase in brain size and the behavioral and cognitive skills [2]. Novel experimental models that are ethically acceptable, user-friendly and recap the human structure have long been sought.

With the advent of innovative stem cell-based 3D tissue models it has been possible to overcome many limitations imposed by animal models or single cell cultures, thus providing the unique opportunity to model human organogenesis. In particular, cerebral organoids are 3D multicellular clusters derived from iPSCs via self-renewal and self-organization and possess certain features of the human brain, including lobes of cortex [3,4]. Moreover, they mimic the cytoarchitecture and the developmental pathways [5] that occur in vivo. Consequently, human cerebral organoids are an excellent experimental model to study neuronal growth, migration and function.

In the early stages of neural development, neuroepithelium is the first structure that appears [6]. Upon exposure to specific stimuli, the neuroepithelial stem cells give rise to the radial glia whose structure is characterized by a ventricular and subventricular zone. Radial glial cells reside in the ventricular zone where they proliferate generating the intermediate progenitors which then migrate to the subventricular zone [3,6]. In mature human cerebral organoids the outer subventricular zone, which is unique to human brain development [3], is composed predominantly by stem cells that complete their neural differentiation mainly in the cortical layer. Moreover, mature human cerebral organoids are characterized by multiple progenitor zones with rosettes, which are apico-basally polarized pseudostratified structures resembling the embryonic neural tube [3,6,7].

Brain development and function require an effective BBB, a dynamic interface between the blood and the brain with the crucial role of maintaining brain homeostasis [8]. Indeed, the BBB strictly controls the passage of blood-borne substances into the central nervous system, and regulates the transport of large and small molecules back into the blood, thereby maintaining the internal milieu which is mandatory for the synaptic activity and neural function [9,10]. The barrier function is granted by the highly specialized characteristics of brain endothelial cells [11], which acquire a unique phenotype during barrier-genesis [12]. Relevant for the maintenance of the BBB is their high levels of junction molecules such as Zonula Occludens-1 (ZO-1) and Vascular Endothelial (VE)-cadherin, components of the tight and adherens junction, respectively. Remarkably, brain endothelial cells release huge amounts of the Brain Derived Neurotrophic Factor (BDNF), to the point that about 50% of the neurotrophin measured in rat brain homogenates coincides with BDNF expressed by cerebral endothelium [11]. Brain endothelial-released BDNF can then interact with its cognate receptor expressed by astrocytes and neurons and exert its protective role against hypoxia, oxidant species and metabolite deprivation [11,13]. Accordingly, some studies indicate that cognitive impairment is associated with decreased BDNF in the endothelium of brain microvasculature [11,14].

Considering the relevant roles of the BBB and, in particular its trophic effect, we here present an experimental model in which human iPSCs-derived cerebral organoids are cultured under an in vitro generated BBB. We compare the structure of cerebral organoids in the presence or not of an in vitro model of BBB.

Whereas it is well known that the endothelium has a major role in restricting the permeability of the BBB, the importance of different cell types -astrocytes and pericytes-in regulating the dynamics of BBB response to physiological or pathological challenges is now appreciated [15]. Moreover, it is known that the neurovascular microenvironment, which includes neural progenitor cells, pericytes, astrocytes, and neurons, contributes to the BBB phenotype. In this work, we utilize a model of BBB, consisting of co-cultured primary capillary brain endothelial cells (HBMEC) and astrocytes in a transwell system [16]. Although simple, this model suffices to detect a higher cortical organization and spatial distribution in BBB-organoids than organoids alone.

## 2. Results

### 2.1 Generation of an in vitro BBB

Since different types of endothelial cells have been used to generate in vitro BBB, we initially analyzed the features of BBB by co-culturing human astrocytes (HA) with either human endothelial cells from the umbilical vein (HUVEC) [17] or capillary brain endothelial cells (HBMEC) [16] using the transwell system. The permeability of the BBB was assessed by adding bovine serum albumin-fluorescein isothiocyanate conjugate (FITC-BSA) (1 mg/mL) to the upper chamber. After 72 h of co-culture astrocytes/endothelial cells, the BBB generated using HBMEC is significantly less permeable than the one generated with HUVEC (Figure 1a, left). Trans-endothelial electrical resistance (TEER) confirmed this result (Figure 1a, right). This difference is linked to the higher amounts of VE-cadherin and ZO-1 in HBMEC than in HUVEC as shown by western blot (Figure 1b). Moreover, immunofluorescence highlights that HBMEC exhibit a more continuous and linear distribution of VE-cadherin and ZO-1 than HUVEC (Figure 1c). Of note, HBMEC synthesize (left panel) and release (right panel) more BDNF than HUVEC as measured by ELISA (Figure 1d). These results prompted us to utilize HBMEC in all the following experiments. This BBB model is rather simple and has previously yielded interesting results by modulating the response of cerebral organoids to different magnesium salts [16]. More sophisticated BBB model can be generated, and will be target of future studies.

**Figure 1.**
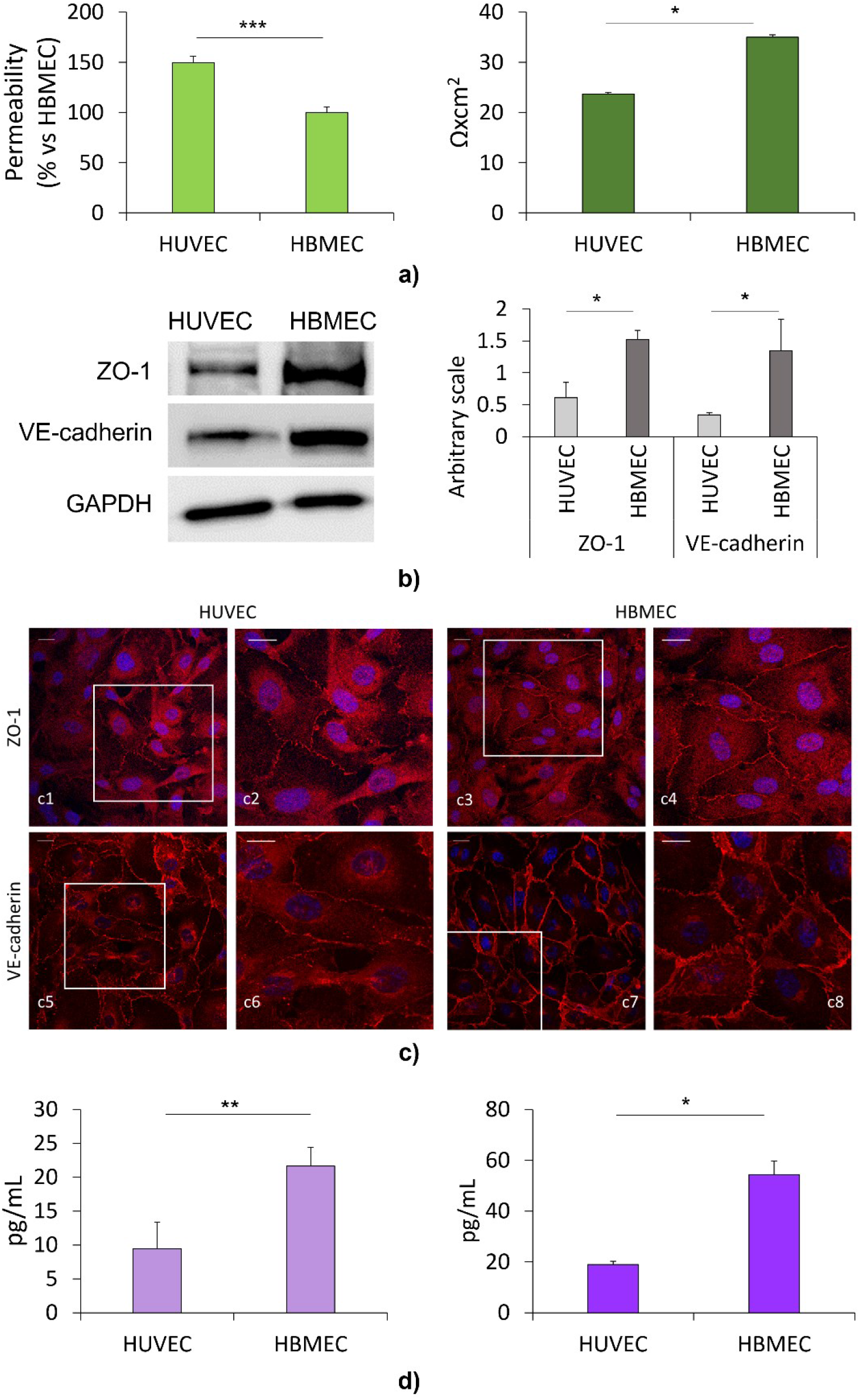
Comparison of BBB generated by co-culturing HA with either HUVEC or HBMEC. a) BBB permeability (left) and TEER (right). The graphs show the quantification of three independent experiments in triplicate. b) Western blots on HUVEC and HBMEC lysates were performed using antibodies against ZO-1 and VE-cadherin. GAPDH was used as a control of loading. A representative blot is shown (left panel). Densitometry was performed on three independent experiments (right panel). c) Immunofluorescence on HUVEC and HBMEC was performed using antibodies against ZO-1 and VE-cadherin (red). DAPI (blue) labels the nuclei. Boxed regions in c1,c3,c5 and c7 are zoomed in c2,c4,c6 and c8 respectively. Scale bar: 15 μm. d) Intracellular (left panel) and released (right panel) BDNF was measured by ELISA assay. The graphs show the quantification of three independent experiments in triplicate. Data are shown as the mean ± standard deviation. *\* p* ≤ 0.05, *\*\* p* ≤ 0.01 and *\*\*\* p* ≤ 0.001.

### 2.2 Evaluation of BDNF amounts in cerebral organoids cultured in the presence or absence of the BBB

It is known that BDNF has a critical role in controlling neurodevelopment, neuroplasticity and neurosurvival. For this purpose, we assessed the expression of the neurotrophin in controls and BBB-organoids. Real time PCR shows that BDNF transcript does not change in the organoids in the presence or absence of the BBB, while ELISA detects higher amounts of the protein in BBB-organoids than in organoids alone (ORG) (Figure 2a and b). We investigated whether the BBB represents a source of BDNF in our experimental model. We anticipate that the high amounts of BDNF released in the media in the presence of the BBB might be due to the contribution of astrocytes and/or HBMEC. Hence, by ELISA we evaluated the amount of BDNF released in the media by i) human cerebral organoids (ORG) ii) human cerebral organoids cultured for 4 days under the BBB (BBB+ORG) iii) the BBB alone iv) human astrocytes (HA) or v) HBMEC. Figure 2c shows that soluble BDNF is significantly higher in the media of BBB-organoids than organoids alone. Our results indicate that the BBB release very high amounts of BDNF, and that HBMEC are the principal source of BDNF in our experimental model.

**Figure 2.**
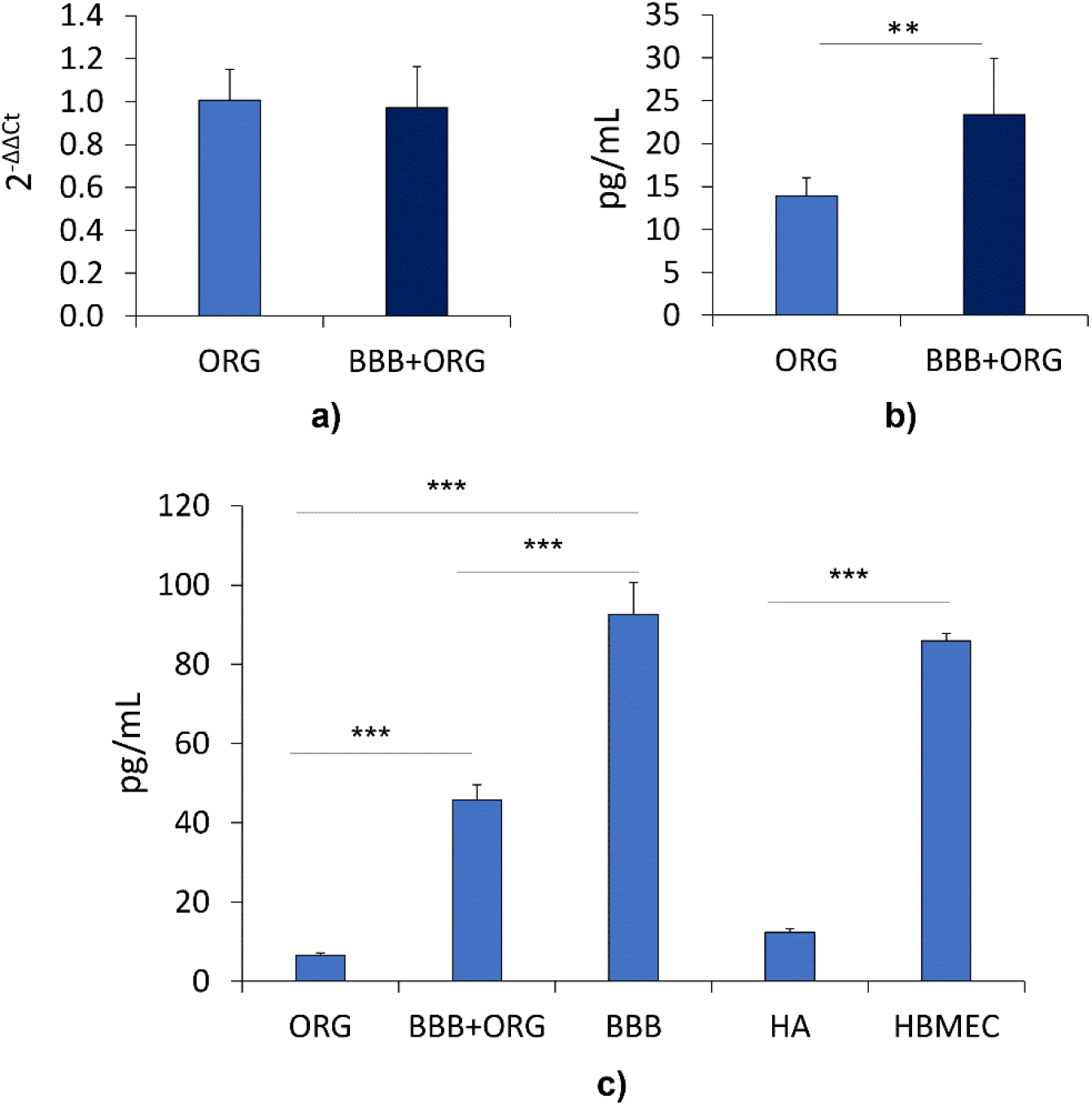
Evaluation of BDNF levels in the absence (ORG) or presence (BBB+ORG) of the BBB. a) *BDNF* expression in cerebral organoids was analyzed by Real time PCR. **b)** BDNF was measured in the human cerebral organoids by ELISA. **c)** Evaluation of BDNF released in the culture media by cerebral organoids alone (ORG), cerebral organoids co-cultured with the BBB (BBB+ORG), the BBB alone (BBB), human astrocytes (HA) and HBMEC alone (HBMEC) by ELISA. The graphs show the quantification of three independent experiments. Data are shown as the mean ± standard deviation. ** *p* ≤ 0.01, *** *p* ≤ 0.001.

### 2.3 Structure of cerebral organoids cultured in the presence or absence of the BBB

Based on previous studies [16], embryoid bodies (EBs) generated from iPSCs were induced to neuroectodermal fate. After 35 days, when a complex morphology with heterogeneous regions containing neurons and astrocytes can be observed [16], cerebral organoids were cultured in the presence (BBB+ORG) or in the absence (ORG) of the BBB for additional 4 days. Sections were obtained and stained with antibodies against γ-Aminobutyric Acid B Receptor (GABA_B_-R) to visualize neurons and SRY-box transcription factor (SOX)2 to detect neuroprogenitors. After staining with anti GABA_B_-R antibodies, confocal microscopy highlights an accumulation of neurites in the marginal zone of BBB-organoids (BBB+ORG) (Figure 3, right panel, green arrows), suggesting a better cortical organization than in controls (ORG) (Figure 3). Of note, in BBB-organoids SOX2-positive cells are observed in the ventricular zone-like layer (white arrows), while in controls the neural-progenitors are also visible in the marginal zone (white arrows), indicating a reduction of the initial lamination.

**Figure 3.**
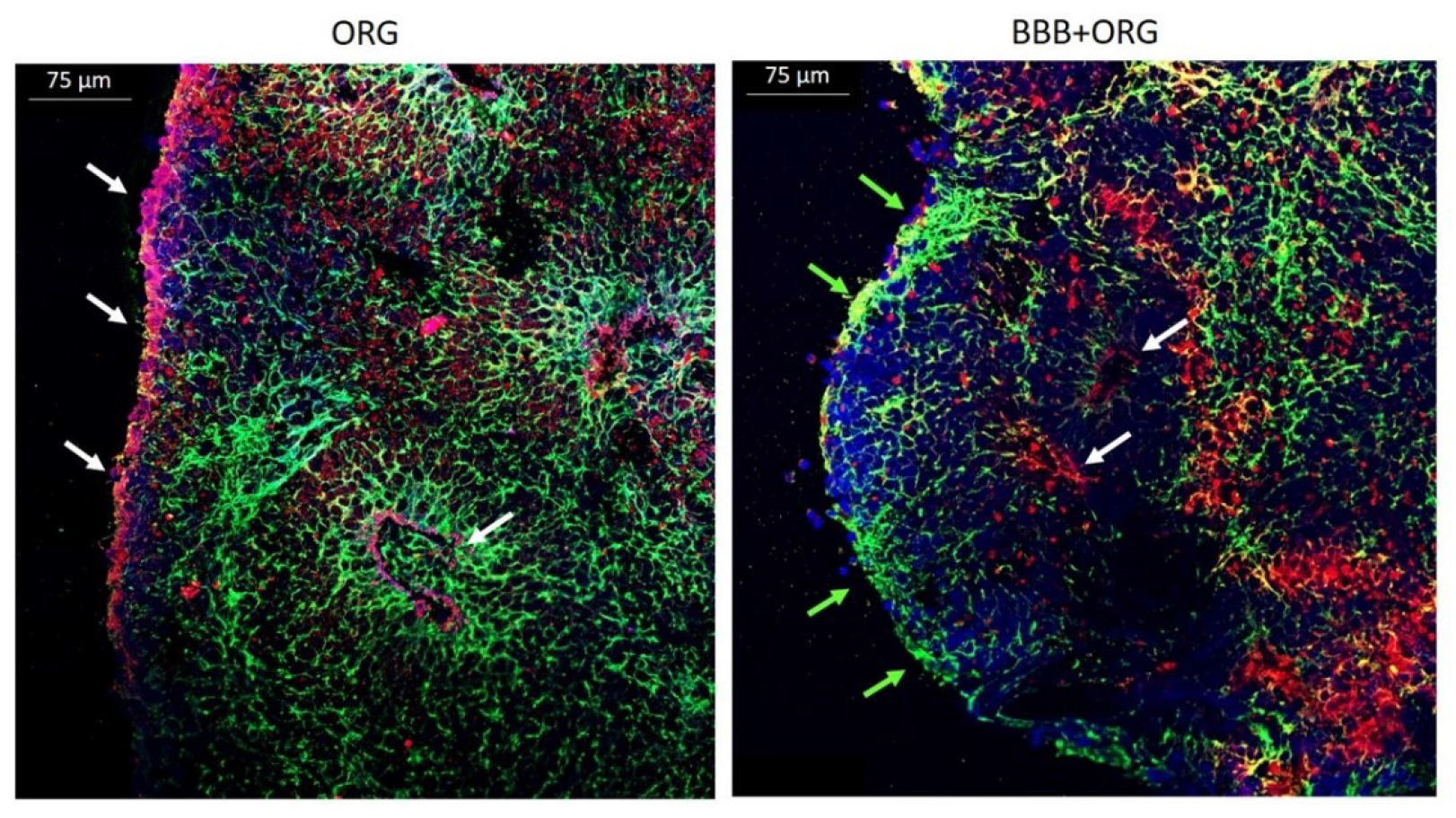
Immunofluorescence staining of a cerebral organoid in the presence (BBB+ORG) or absence (ORG) of the BBB. Antibodies against the neuronal marker GABA_B_-R (green) and the neural progenitor marker SOX2 (red) were utilized. DAPI (blue) labels the nuclei. For lower magnification please see Supplementary (Figure S1).

On the basis of these findings, we analyzed the structure of cerebral organoids cultured in the presence or absence of the in vitro generated BBB for 4 days using optical microscopy (Figure 4, a1-b1) and transmission electron microscopy (TEM) (Figure 4, a2-7, b2-7) on resin embedded organoids fixed with glutaraldehyde and osmium tetroxide.

**Figure 4.**
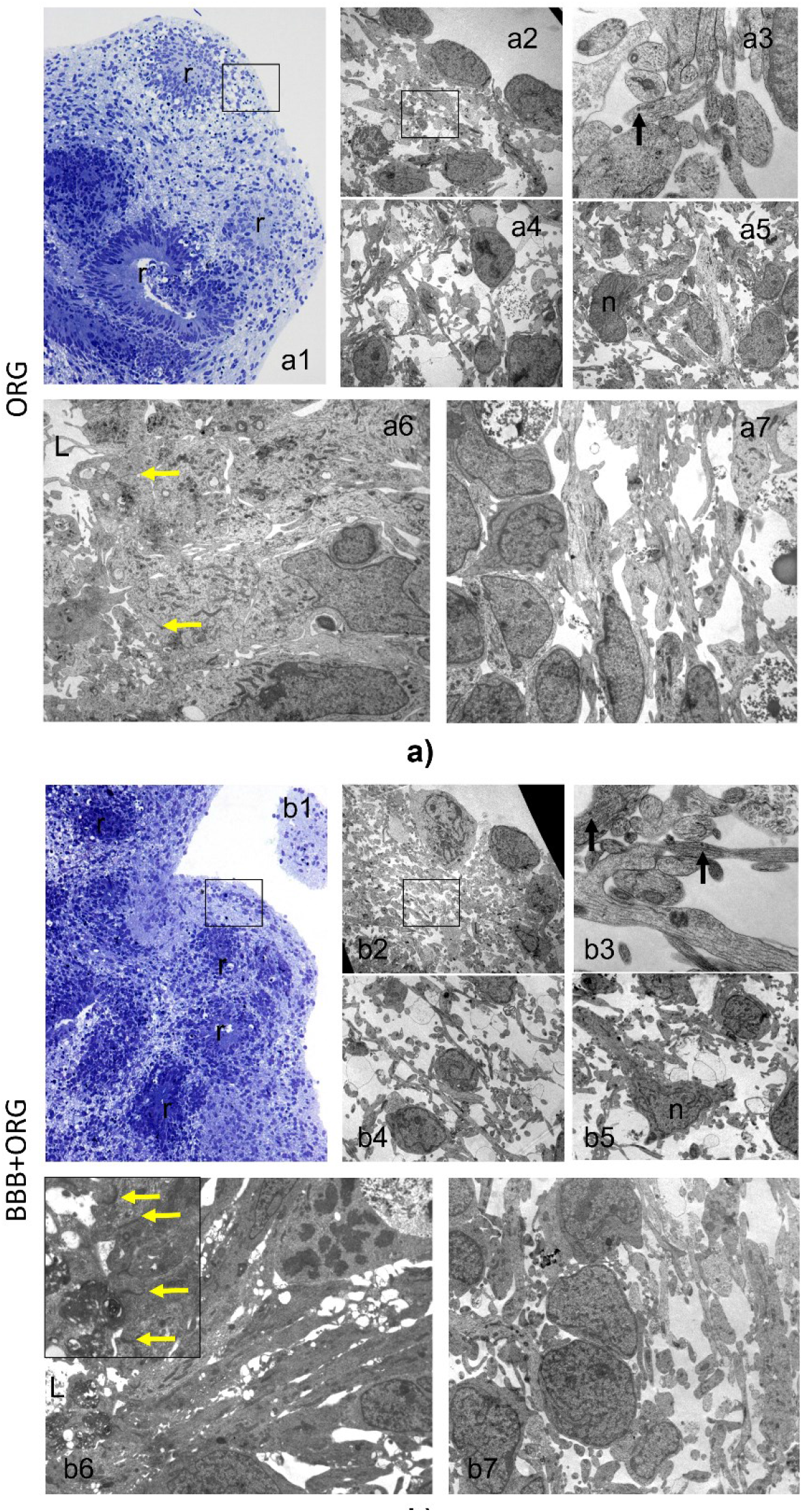
Structure of a cerebral organoid in the presence (b) or absence (a) of the BBB. Semithin resin sections were stained with toluidine blue and analyzed at light microscope, then thin sections from selected areas (boxed regions) were analyzed by TEM. a1, b1: toluidine blue stained semithin sections i.o. x200; a2-5 and b2-5: electron micrographs, original magnification: a2, a4, a5, b2, b4×3000; b5×4400; a3, b3 x12000; r rosette, n neuron like cell. Black arrows indicate neurites. Ultrastructure of neuroepithelial rosettes in cerebral organoid in the presence (b6 and 7) or absence (a6 and 7) of the BBB. Original magnification a6, b7: x4400; a7, b6: x3000 x7000. L= neural tube-like lumens. Yellow arrows indicate epithelial junctions.

By optical microscopy, toluidine blue staining confirms differences between organoids co-cultured or not with the BBB. In BBB-organoids (Figure 4, insert b1) neuroepithelial rosettes are smaller than in the controls without the BBB (Figure 4, insert a1) [18– 20]. Of note, the cortical zone of BBB-organoids is characterized by cells with a better spatial organization than the control (Figure 4, inserts b1 and a1, boxed regions). Moreover, an initial lamination and cortical multilayer formation can be observed.

At the periphery of the neuroepithelial rosettes, different cell types in organoids and BBB-organoids are detected by ultrastructural studies. Figure 4 shows neural-like cells [21,22] with a voluminous nucleus and axon-like extensions (Figure 4, “n” in inserts a5 and b5), and round cells with scarce cytoplasm and no clear extension, usually coupled with neurites (Figure 4, inserts a4, b4), a morphology which is indicative of a glial phenotype. In addition, groups of neurites with microtubules and core-dense vesicles can be observed (Figure 4, boxed regions in inserts a2 and b2, enlarged views in a3 and b3). At the periphery, cerebral organoids are delimited by cells with short neurites-like extensions inward (Figure 4, inserts a2 and b2). We then focused on the ultrastructural features of the neuroepithelial rosettes. By TEM, no major differences emerge in the organization of the rosettes between organoids and BBB-organoids. Indeed, all the rosettes are surrounded by neurites and cells corresponding to the deeper layers of the developing cortex. Moreover, in both samples well-developed epithelial junctions are visible (Figure 4, arrows in inserts a6 and b6, boxed region in insert b6) delimiting neural tube-like lumens (Figure 4,”L” on the left in inserts a6 and b6).

## 3. Discussion

Cerebral organoids have yielded unprecedented progresses in our understanding of neurodevelopment as well as of some aspects of neuropathology. The weakness of this in vitro system is the lack of the endothelial component, which plays fundamental roles in brain homeostasis and also in brain development [23]. Various experimental approaches have been utilized to vascularize brain organoids. Some vessels formed but no perfusion occurred upon iPSCs reprogramming into endothelial cells [24]. Genetically engineered embryonic stem cells originated a vascular network which remained mainly confined on the surface of the organoids. Interestingly, the endothelial cells expressed BBB markers, such as ZO-1, and their presence accelerated the functional maturation of the organoids [25], thus confirming the role of endothelial cells and BBB in brain formation and function [26]. Indeed, cerebral endothelial cells, neuronal and glial progenitors communicate by exchanging signals that model proliferation and differentiation [27].

In order to have cost effective and user friendly experimental models, we propose to culture human cerebral organoids in the presence of a rather simple in vitro generated BBB. We have previously developed BBB using human astrocytes and HUVEC [17]. However, significant differences were highlighted between HUVEC and HBMEC, which upregulate factors actively involved in neuroprotection and cell differentiation, angiogenesis, and immunoregulation [28]. Therefore, in this paper we utilize HBMEC, which show higher amounts and a more continuous and linear distribution of junctional proteins than HUVEC, thus granting the generation of a highly impermeable BBB. These cells also secrete relevant amounts of BDNF, which promotes the differentiation of cortical progenitor cells and then their maturation into neurons [29]. It is known that adding BDNF to the medium optimizes neural induction in human organoids [30], thus offering an explanation to the aforementioned results reported by Cakir et al [25]. This consideration prompted us to evaluate the structure of cerebral organoids cultured or not under the BBB generated using HBMEC. In BBB-organoids the cells in the cortical zone are spatially organized, with the presence of areas of initial lamination and cortical multilayer formation. In parallel, ultrastructural studies confirm the higher cortical organization and better spatial distribution in BBB-organoids than organoids alone, with alternating arranged layers of neurites’ bundles and cells. These aspects are evocative of a more advanced stage of differentiation in BBB-organoids [20].

We therefore reasoned that the more mature phenotype of BBB-organoids might be linked to the synthesis and release of BDNF by the BBB. Indeed, BDNF plays an important regulatory role from the earliest steps of developing brain and acts as a differentiation factor for most neurons in the central nervous system [31]. In our experimental model, HBMEC are the principal source of released BDNF. We also show that the levels of BDNF are higher in BBB-organoids than in organoids alone. This last result is intriguing since in embryonic neurons BDNF acts as an autocrine factor in stimulating axon formation by the rapid activation of different signal transduction pathways [32]. Moreover, since BDNF is positively charged, upon secretion it immediately binds to the cell surface or to components of the extracellular matrix nearby [33]. More experiments are necessary to identify the intra or extracellular localization of BDNF in our model. Another question raised by our results is whether the levels of cerebral BDNF are modulated during neurodevelopment. At the moment it is known that BDNF expression increases with the maturation of the central nervous system in developing rats [34].

In BBB-organoids BDNF transcript is not modulated. Therefore, we hypothesize that post transcriptional and/or post-translational mechanisms are implicated in the upregulation of BDNF. It is also feasible that BDNF diffuses in the organoids from the extracellular environment, an event that might explain why there is less BDNF detected in the medium of BBB-organoids than BBB alone. Alternatively, soluble BDNF might be the target of extracellular proteases released by the organoids or by the cellular components of the BBB.

We are aware of the limitations of this study. First, all the experiments were performed on organoids after 35 days of culture. It would be interesting to maintain organoids at different stages of maturation under the BBB for different times. Second, we focused on BDNF because it shows a more widespread distribution than other neurotrophins including Nerve Growth Factor [35]. However, it is likely that other neurotrophins and growth factors are involved. Third, it would be challenging to investigate the role of BDNF, for instance by silencing BDNF in cerebral endothelial cells. While more experiments are required to better characterize and improve our experimental setting, we propose that it represents a rather simple model that takes the role of BBB into consideration, thus being useful not only for studying neurogenesis but also for drug screening.

## 4. Materials and Methods

### 4.1 Generation of human cerebral organoids

Human cerebral organoids are generated from human induced pluripotent stem cells (iPSCs) which were purchased from GibcoTM (Gibco by Thermofisher, Waltham, Massachusetts, USA). iPSCs were plated in a dish pre-coated with Matrigel® hESC-Qualified Matrix (Corning, Corning, New York, USA) and cultured in mTesR culture media (Stem Cell Technologies, Vancouver, Canada) following the manufacturers’ instructions. The cells were routinely tested for the expression of stemness markers and used for 4–6 passages. To generate cerebral organoids, iPSCs are detached using a gentle cell dissociation reagent (Stem Cell Technologies) once they reach the 80% of confluence as indicated in the Lancaster protocol [3]. In brief, to generate embryoid bodies (EB), 9000 cells/well are plated in ultra-low attachment 96-wells plate (Corning) and maintained in media containing 4ng/mL Basic Fibroblast Growth Factor (bFGF) (Thermo Fisher Scientific, Waltham, Massachusetts, USA), which maintains the pluripotency of iPSCs [3], and 10 µM Rock-inhibitor Y-27632 (Sigma-Aldrich, St. Louis, Missouri, USA) to increase cell survival. Then, EB reach about 500 µm in diameter and are transferred into a neural induction medium for 4 days. At this point, neuroepithelia can be visualized. Then, the neuroectoderm formations are embedded in the centre of a droplet of Matrigel® hESC-Qualified Matrix (Corning, Corning, New York, USA) and cultured for other 4 days in a differentiation medium without Vitamin A. At this stage, the tissues become more complex with some budding outgrowth and radial processes and the immature organoids are maintained on an orbital shaker to favour nutrients and oxygen exchange. Finally, organoids are transferred into a new differentiation medium added with vitamin A for the last twenty days. At the end of these steps, human cerebral organoids are mature and ready for the experiments. The cerebral organoids are maintained in culture for additional 4 days in the presence (BBB+ORG) or absence (ORG) of an in vitro generated BBB. To perform all the analysis, a pool of two organoids was used for each experimental replicate.

### 4.2 In vitro model of blood brain barrier (BBB)

Human brain microvascular endothelial cells (HBMEC) (Innoprot, Bizkaia, Spain) were cultured in ECM medium supplemented with 5% of fetal bovine serum (FBS), 1% of endothelial cell growth supplement (ECGS) and 1% of penicillin/streptomycin solution (P/S) (Innoprot). Human umbilical vein endothelial cells (HUVEC) were obtained from the American Type Culture Collection (ATCC, Manassas, WV, USA) and cultured in medium M199 (Euroclone, Milan, Italy) containing 10% fetal bovine serum (FBS), 1 mM L-Glutamine, 1mM Sodium Pyruvate, 1 mM Penicillin-Streptomycin, 5 U/mL Heparin and 150 μg/mL Endothelial Cell Growth Factor on 2% gelatin-coated dishes (Euroclone). Human astrocytes (HA) (Science Cell, Carlsbad, USA) were cultured in HA medium containing 2% of FBS, 1% of astrocyte growth supplement and 1% of P/S. For the experiments, all the cells were used between 4-6 passages. The BBB in vitro model is generated exploiting the Transwell system (Corning) with a 0.4 µm pore size. At first, human astrocytes are seeded (35,000/cm^2^) on the underside of the transwell insert precoated with Poly-L-lysine (2 µg/mL) (Sigma-Aldrich). Then, HBMEC or HUVEC are seeded on the upper side (60,000/cm^2^) precoated with fibronectin (50 µg/mL) (Sigma-Aldrich). 72 h after the co-culture, the BBB reaches the maximal resistance and it is ready to perform the experiments [17].

### 4.3 Permeability assay and TEER

To validate the BBB model, a permeability assay and trans-endothelial electrical resistance (TEER) were performed 72 h after the co-culture of HA and endothelial cells. The permeability was measured exploiting the bovine serum albumin-fluorescein isothiocyanate conjugate (FITC-BSA) (Sigma, Saint Louis, Missouri, USA) [36]. Briefly, FITC-BSA (1 mg/mL) was added to the upper chamber and, after different times, the medium in the lower chamber was collected. By using the Varioskan LUX Multimode Microplate Reader (Thermo Fisher Scientific), the fluorescence was measured at excitation/emission wavelength of 495/519 nm. All the experiments were repeated 3 times in triplicates. Results are shown as % of HUVEC vs HBMEC. To measure the TEER of the endothelial monolayers, an EndOhm (World Precision Instruments, Friedberg, Germany) was used. The experiments were repeated three times in triplicates. Results were expressed in Ωxcm^2^ [37], and presented as the mean ± standard deviation (SD).

### 4.4 Western blot

The cells were lysed in 50 mM Tris–HCl (pH 7.4) containing 150 mM NaCl, 1% NP40, 0.25% sodium deoxycholate, protease inhibitors (10 μg/mL Leupeptin, 10 μg/mL Aprotinin, 1 mM PMSF) and phosphatase inhibitors (1 mM sodium fluoride, 1 mM sodium vanadate, 5 mM sodium phosphate). By using the Bradford reagent (Sigma Aldrich), the protein concentration was assessed. Lysates (30 μg/lane) were separated on SDS–PAGE and transferred to nitrocellulose sheets using the Trans-Blot Turbo Transfer System following the manufacturer’s instructions (Bio-Rad). The immunoblot analysis was performed using antibodies against ZO-1 (Thermo Fisher Scientific), VE-cadherin (Cell Signaling Technology, Danvers, Massachusetts, USA) and Glyceraldehyde-3-Phosphate Dehydrogenase (GAPDH) (Santa Cruz Biotechnology, Dallas, Texas, USA), which was used as control of loading. Then, the nitrocellulose membrane was washed and incubated with secondary antibodies labeled with horseradish peroxidase (Amersham Pharmacia Biotech Italia, Cologno Monzese, Italy). Immunoreactive proteins were detected with ClarityTM Western ECL substrate (Bio-Rad Laboratories, Hercules, CA, USA). Each experiment was performed three times. The Image J software (Version 1.52a, National Institutes of Health, Bethesda, MD, USA) was utilized to perform the densitometric analysis. A representative blot is shown, and the graph is obtained by the measurements of three independent experiments ± SD.

### 4.5 Fluorescence Microscopy

After fixation in phosphate buffered saline containing 4% paraformaldehyde and 2% sucrose (pH 7.6), HUVEC and HBMEC were incubated with the antibodies against VE-cadherin (Cell signaling) or ZO-1 (Thermo Fisher Scientific) and processed for immunofluorescence. The human organoids were frozen in Optimal Cutting Temperature compound (OCT) and cryosectioned at 10 µm. The sections were collected on coverslips and stained with anti-GABA_B_-R to detect neurons [38], anti-SRY-box transcription factor (SOX)2 to visualize neural progenitors [39]. The goat anti-mouse Alexa Fluor 488 (green) and the goat anti-rabbit 546 (red) were used as secondary antibodies. 4′,6-diamidino-2-phenylindole (DAPI) was used to stain the nuclei (blue). Finally, the slices were mounted with moviol. Images were acquired using a 40X objective in oil by an SP8 confocal microscope (Leica Microsystems, Buffalo Grove, IL, USA).

### 4.6 Light and transmission electron microscopy -TEM-on resin embedded organoids

Cerebral organoids cultured in the presence or absence of the in vitro generated BBB for 4 days were fixed in 2,5% buffered glutaraldehyde, post-fixed in 1% Osmium tetroxide buffered solution, and embedded in epoxy resin (Embed 812 kit for Electron Microscopy, EMS Company) using standard preparative procedure for ultrastructural analyses of tissues and cells. Semithin resin sections were stained with toluidine blue and analyzed at light microscope. Then, thin sections from selected areas were stained with uranyl acetate/lead citrate and examined by EM-109 ZEISS and CCD-Megaview G2 (I-TEM imaging platform software). Image of each organoid was reconstructed using Leica LAS X Navigator Software.

### 4.7 Real time PCR

Total RNA from cerebral organoids was extracted using the Purelink-RNA Mini kit (Invitrogen, Waltham, Massachusetts, USA) Single-stranded cDNA was synthesized from 0.6 µg RNA in a 20 µL final volume using the High Capacity cDNA Reverse Transcription Kit, with RNase inhibitor according to the manufacturer’s instructions (Applied biosystems, Waltham, Massachusetts, USA). The Real time PCR analysis was performed three times in triplicate using the CFX96 Real Time PCR Detection system (Biorad, Hercules, California, USA). TaqMan Gene Expression Assays probes were used (Life Technologies, Monza, Italy). The probes were targeting BDNF (Hs02718934_s1) and the housekeeping gene GAPDH (Hs99999905_m1), that was used as an internal reference. Relative changes in the gene expression were analyzed with the 2^−ΔΔCt^ method ± SD.

### 4.8 ELISA assay

To determine the protein amount of human BDNF in our samples, the ELISA kit for human BDNF protein was used according to the manufacturer’s instructions (Abcam, Cambridge, UK) [16]. ELISA was performed on 30 μg of protein extracts or on 50 µL of culture media in triplicates at least three times ± SD.

### 4.9 Statistical Analysis

Data are reported as media ± SD. The data were analyzed using the non-parametric Mann Whitney test. Statistical significance was defined for *p*-value ≤ 0.05. * *p* ≤ 0.05; ** *p* ≤ 0.01; *** *p* ≤ 0.001.

